# Gene-specific reactivation of X-linked genes upon Xist loss is linked to the chromatin states in extraembryonic endoderm and epiblast stem cells

**DOI:** 10.1101/2023.10.20.563299

**Authors:** M Arava, S Majumdar, LS Sowjanya, HC Naik, R Baro, S Gayen

## Abstract

In eutherian mammals, X-chromosome dosage between sexes is balanced through the inactivation of one of the two X-chromosomes in female cells. In mouse, X-inactivation initiates at ∼4-8 cell stages of embryogenesis, where paternal-X undergoes imprinted X-inactivation. Subsequently, it switches to random X-inactivation in post-iplantation epiblast. The initiation of XCI is orchestrated by Xist. However, the role of Xist in the maintenance of X-chromosome inactivation remains underexplored. Here, we have explored the role of Xist in the maintenance of X-inactivation in extraembryonic endoderm stem cells (XEN) and epiblast stem cells (EpiSC), which undergo imprinted and random form of X-inactivation respectively. We show that removal of Xist leads to the partial reactivation of inactive-X chromosome. Intriguingly, many reactivated genes were found to be common between XEN and EpiSC, indicating these genes require Xist to maintain their silent state irrespective of the lineages or forms of X-inactivation. Notably, despite Xist ablation and the subsequent removal of DNA methylation, several X-linked genes remained resistant to reactivation, indicating the involvement of other factors in maintaining the silencing of these genes. On the other hand, we show that genes on the inactive-X with low levels of H3K9me3 and high levels of H3K27me3 are more susceptible to reactivation upon the loss of Xist. Interestingly, active-X homolog of the reactivated genes was found to be enriched with H3K4me3 and H3K27ac. Taken together, our study sheds light on the role of chromatin states in the reactivation of X-linked genes following the loss of Xist in XEN and EpiSC.

## Introduction

In eutherian mammals, for balancing the X-linked gene dosage between sexes, one X-chromosome in female get transcriptionally silenced through the process known as X-chromosome inactivation (XCI) (1). In mouse, there are two forms of X-inactivation: imprinted XCI (I-XCI) and random XCI (R-XCI) (2, 3). The I-XCI initiates at the two – four cell stage of the developing embryo, where paternal X preferentially gets inactivated (4). Subsequently, the inactivated-X gets reactivated in the inner cell mass (ICM) of the late blastocyst and undergo R-XCI in the embryonic epiblast: i.e., either paternal or maternal X chosen randomly for inactivation (5–8). In recent years, the factors responsible for the initiation and establishment of X-inactivation have been extensively studied. Initiation of XCI occurs through a series of ordered epigenetic events. Importantly, XCI are prefaced by the expression of the long non-coding RNA (lncRNA) Xist from the prospective inactive-X and followed by recruitment of different chromatin modifiers such as PRC1, PRC2, SPEN, RBM15 etc. (9, 10). Subsequently, CpG methylation stabilize the silenced state (11). However, the factors and mechanisms governing the maintenance of X-inactivation remains poorly understood. Understanding of mechanisms involve in maintenance of X-inactivation is crucial as failure to do so can result in a wide array of abnormalities, including cellular lethality.

Xist is required for the initiation of both imprinted and random X-inactivation. The deletion of paternal Xist in the developing mouse embryo results in embryonic lethality due to the defects in I-XCI (12). Similarly, the loss of Xist at the onset of initiation of R-XCI leads to defects in dosage compensation (13). Interestingly, the heterozygous loss of Xist leads to non-random X-inactivation towards WT allele (14). However, the role of Xist in the maintenance of X-inactivation remains poorly understood. Previously it was thought that the Xist is dispensable for the maintenance of XCI as loss of Xist did not lead to the reactivation of the inactive-X (15–18). However, most of these studies assayed inactivation status of the X-chromosome based on few X-linked genes or reporter genes. Later on, studies involving chromosome wide analysis of X-linked gene expression or more sensitive assays revealed that loss of Xist leads to partial reactivation of X-linked genes, suggesting that Xist is not completely dispensable for the maintenance of XCI (19, 20, 29, 30, 21–28). Notably, the degree of reactivation varies among different cell types and the loss of Xist in different tissues leads to diverse outcomes, suggesting tissue-specific role of Xist in the maintenance of XCI (22, 27, 29). However, the mechanism underlying the variable effects of Xist loss among X-linked genes and different tissues remain unknown. Moreover, most of the above-described studies pertain to somatic lineages, which undergo random X-inactivation, therefore role of Xist in the maintenance of imprinted X-inactivation remains unknown.

In this study, we have investigated the role of Xist in the maintenance of imprinted and random XCI in early embryonic lineages: extra embryonic endoderm cells (XEN) and epiblast stem cells (EpiSC). These cells represent a developmental window, where they transit to the initiation to the maintenance phase of XCI. Notably, XEN cells represent the primitive endoderm of blastocyst and undergo I-XCI, whereas EpiSC represents the embryonic epiblast of post-implantation embryo undergo R-XCI. We show that loss of Xist in XEN and EpiSC leads to the partial reactivation of the inactive-X, suggesting many genes can maintain their silent state even without Xist. Surprisingly, we found that pool of genes reactivated were common between the two lineages, indicating that these genes are dependent on Xist to maintain their silent state in both EpiSC and XEN cells. Furthermore, we have explored the role of CpG DNA methylation in the maintenance of imprinted inactive-X in XEN cells. Finally, our research revealed that chromatin states of X-linked genes play a decisive role towards susceptibility to the reactivation of the X-linked genes following the loss of Xist in XEN and EpiSC. Taken together, our study provides significant insight into the mechanisms governing the maintenance of imprinted and random XCI within early embryonic lineages.

## Results

### Ablation of Xist in XEN cells using CRISPR-Cas9

It is believed that Xist lncRNA plays important role in the initiation of imprinted X-chromosome inactivation. However, its contribution to the maintenance of imprinted X-chromosome inactivation remains unknown. It is known that Xist continues to be highly enriched to the inactive-X even after the initiation and establishment of the XCI. Therefore, we investigated whether the Xist loss from the inactive X would perturb the maintenance of imprinted XCI. To investigate the role of Xist in the maintenance of the imprinted XCI, we selected XEN cells as our model system (Fig. 1A). XEN cells are stem cells, which represents the primitive endoderm cells of the blastocyst and stably propagate the inactivated paternal X-chromosome and therefore serves as a model system for studying the maintenance phase of the imprinted XCI (Fig. 1A). Notably, these XEN cells are hybrid, carrying polymorphic X-chromosomes (X^Mus^: *Mus musculus* origin and X^Mol^: *Mus molossinus* origin) and thereby allowed us to differentiate the X-linked gene expression between active-X vs. inactive-X through conducting allele-specific expression analysis of X-linked genes (Fig. 1A). To ablate Xist expression in XEN cells, we used CRISPR-Cas9 approaches. We targeted Xist promoter including Xist exon 1 using a pair of small guide RNAs (sgRNA) as described earlier (31) (Fig. 1B). We screened puromycin resistant single cell derived clones through polymerase chain reaction (PCR) and gel electrophoresis, followed by Sanger sequencing. Using this approach, in our previous study we generated one △Xist-XEN clone (designated here as △Xist-Pat-3) with heterozygous paternal Xist deletion (31). In this study, we created two additional clones (△Xist-Pat-1 and △Xist-Pat-2) with similar heterozygous paternal Xist deletion (Fig. 1C and 1D). We profiled the the pattern of heterozygous deletion, whether it originated from paternal or maternal X-chromosome, we leveraged the strain-specific single nucleotide polymorphism (SNP) present in the deleted PCR amplicon (Fig. 1E). Moreover, Sanger sequencing of the deleted PCR product confirmed that the sgRNA targeted sites had been joined, affirming the deletion of the targeted segments (Fig. 1D and 1F). Next, we coducted RNA-sequencing (RNA-seq) followed by allele-specific analysis to assess the status of X-linked gene expression from the inactive-X upon Xist deletion. From the RNA-seq data, first we investigated the status of Xist expression in the paternally Xist deleted clones. As anticipated, we found that in WT XEN cells, Xist exclusively expressed from the paternal X-chromosome, thus validating imprinted inactivation of the paternal X-chromosome in these cells (Fig. 2A). In consistence, there was no expression of Xist neither from paternal or maternal allele in all three paternal Xist deleted clones (△Xist-Pat-1, △Xist-Pat-2 and △Xist-Pat-3), suggesting complete loss of Xist expression in these cells (Fig. 2A).

**Figure 1:**
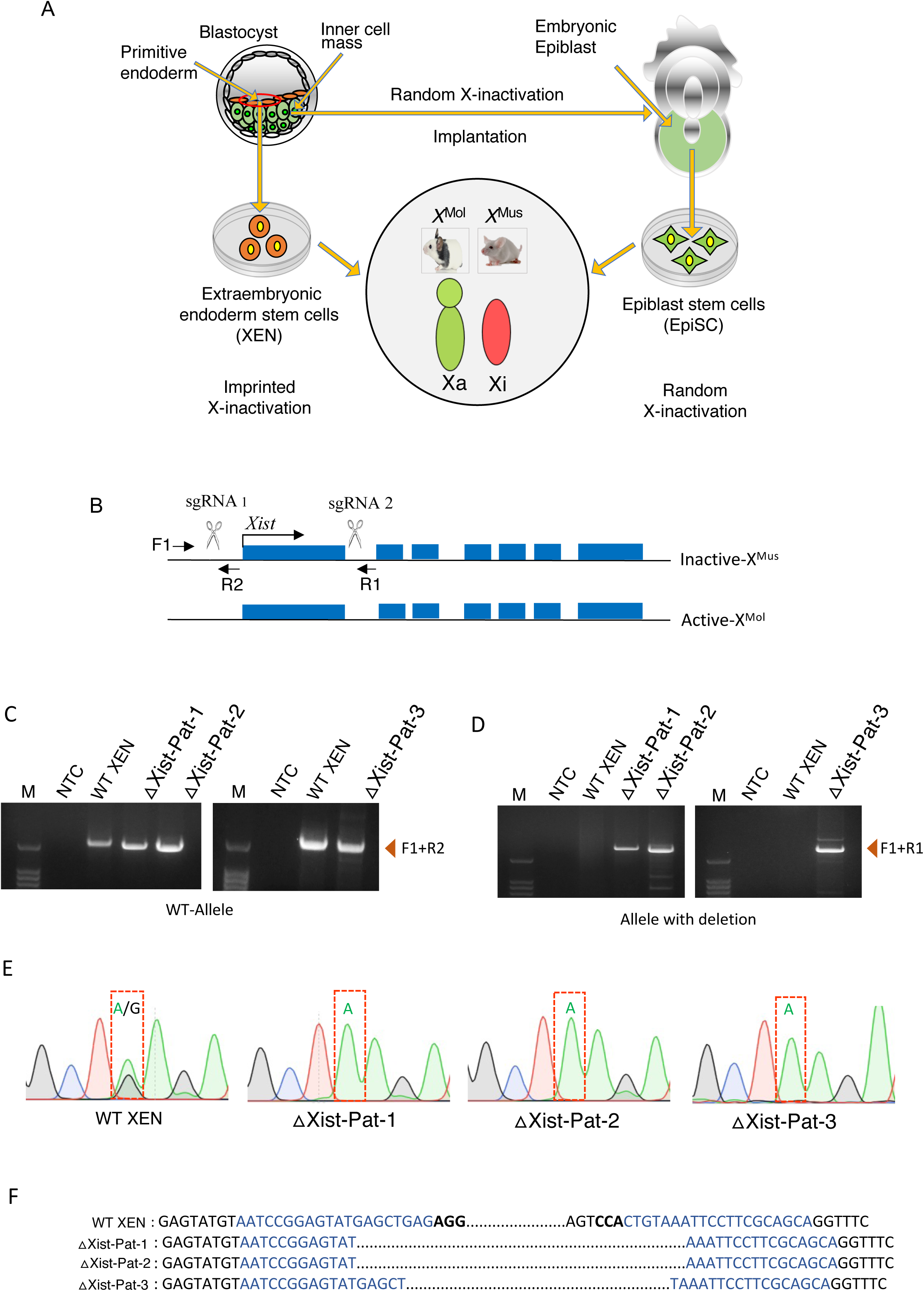
Generation of XEN cell lines with Xist heterozygous deletion in the inactive-X. (A) Schematic showing the embryonic lineages of XEN and EpiSC used for the study. XEN and EpiSC used for the study are hybrid cells and have polymorphic X-chromosome, which facilitated allele-specific analysis of X-linked gene expression from active vs. inactive X chromosome. In both cell types, X-chromosome of *Mus musculus* origin (X^Mus^) is the inactive-X. (B) Schematic showing the sgRNA target sites, location of screening primers for the identification of wild type (F1 and R2), and Xist deleted (F1 and R1) alleles at the Xist locus. (C, D) PCR genotyping for the identification of Xist (C) wild type (F1+R2) or (D) deleted (F1+R1) allele single cell clones. M: marker, NTC: no template control. (E) Chromatogram showing the SNPs corresponding to paternal allele deletion of Xist in the Xist deleted clones. The dotted box showing the SNP present in the deleted amplicon. (F) Representation of deleted region sequences of wild-type and three independent single-cell clones from top to bottom. The black font correspond to the sgRNA flank sequence, the black and bold font is the protospacer motif (PAM) sequence, and the blue font represents the sgRNA sequence.

**Figure 2:**
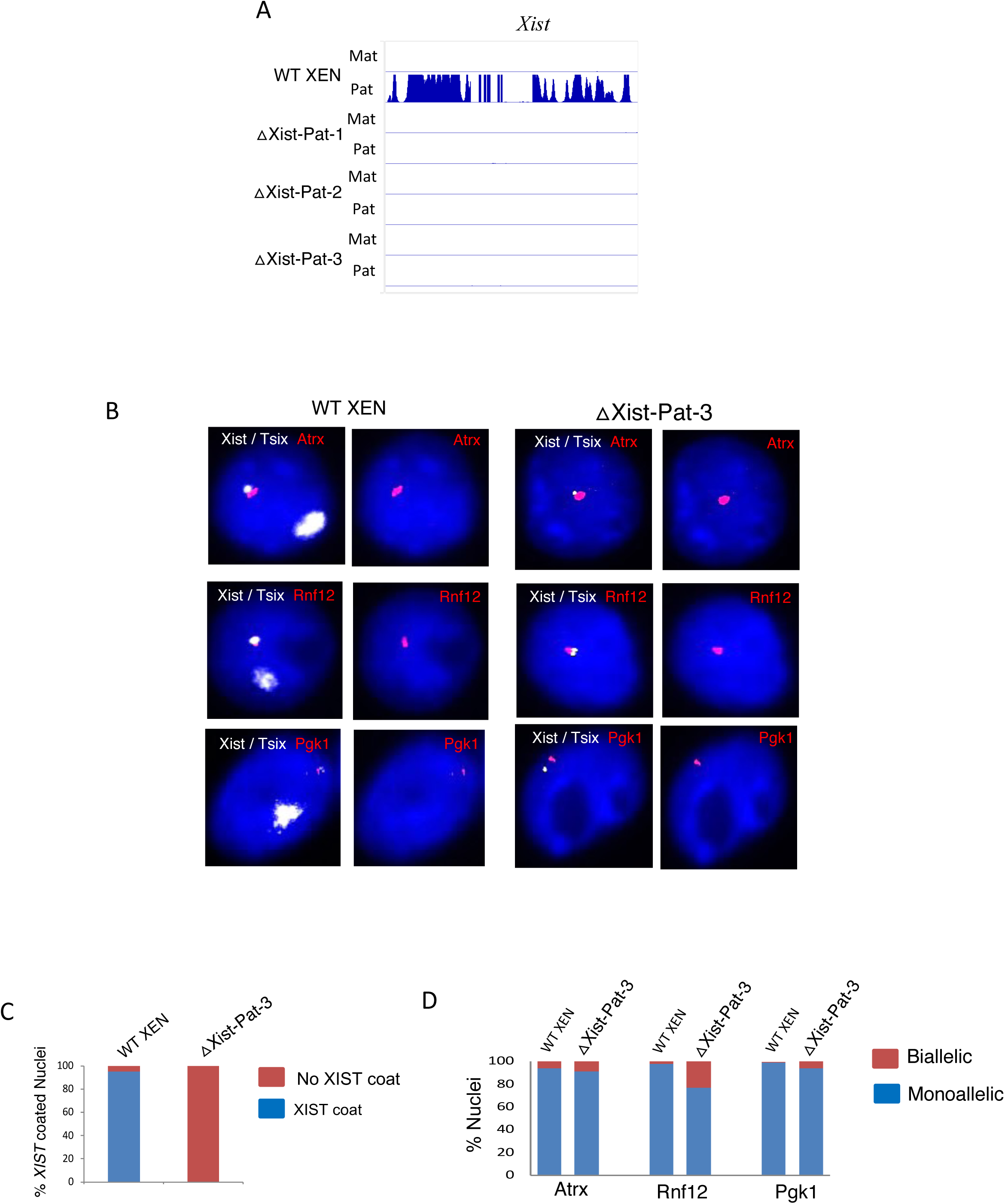
Loss of Xist expression and coating in XEN cells with heterozygous Xist deletion. (A) Genome browser view of the allelic Xist expression in WT and three △Xist clones. (B) Representative images for the RNA-FISH of Xist / Tsix (white color cloud/pinpoint, respectively), and three X-linked genes Atrx, Rnf12, and Pgk1 (detected in red). Nuclei are stained with DAPI (C) Plots representing the quantification of the nuclei with Xist cloud in WT and △Xist clones. (D) Quantification plots for biallelic vs. monoallelic expression of *Atrx*, *Rnf12*, and *Pgk1* in WT and △Xist clones.

### Loss of Xist leads to the partial reactivation of the inactive-X in XEN and EpiSC

Previously, we demonstrated that loss of Xist in △Xist-Pat-3 XEN cells led to the reactivation of a subset of X-linked genes (31). In this study, we have delved deeper into the extent of of reactivation of the inactive-X after the loss of Xist, including other △Xist XEN cells. First, we performed RNA fluorescence in situ hybridization (RNA-FISH) for three X-linked genes: Atrx, Pgk1 and Rnf12 along with the Xist in WT and △Xist-Pat-3 cells (Fig. 2B). As reported earlier, the inactive-X of WT XEN nuclei displayed a crisp, solid, intense Xist coating. In contrast, there was no such coating or expression of Xist in paternally Xist deleted clone (△Xist-Pat-3), confirming the complete loss of Xist expression in these cells (Fig. 2B, 2C). On the other hand, all three X-linked genes we analyzed displayed mono-allelic expression from the active X-chromosome in WT XEN cells (Fig 2B and 2D). Surprisingly, in △Xist XEN cells, all three genes continued to maintain their monoallelic expression despite the loss of Xist, suggesting no reactivation of these X-linked genes (Fig. 2B and 2D). Only Rnf12 showed slight increase in biallelic expression in △Xist XEN cells compared to the WT XEN (Fig. 2B and 2D). To further explore, the expression status of X-linked genes from the inactive-X chromosome in Xist deleted XEN cells, we profiled chromosome wide expression analysis of X-linked genes through allele-specific analysis of RNA-seq data. To profile the reactivation, we calculated fraction allelic expression of X-linked genes from the inactive-X chromosome (Fig. 3A). As anticipated, we found in WT XEN and empty vector transfected XEN cells (WTEV.1, WTEV.2 and WTEV.3), the majority of X-linked genes did not show expression from the inactive-X (Fig. 3A). Only few genes showed biallelic expression, most of which are known to escape XCI (Fig. 3A). Surprisingly, we found that many X-linked genes could maintain their silent state despite the lack of Xist in all three paternal Xist deleted clones (△Xist-Pat-1, △Xist-Pat-2 and △Xist-Pat-3), suggesting these genes do not rely on Xist to maintain the inactive state (Fig. 3A and 3B). Nevertheless, several genes showed expression from the inactive-X chromosome in paternal Xist deleted cells, indicating their dependency on Xist to maintain their silent state (Fig. 3A and 3B). Notably, the extent of reactivation varied significantly among the paternal Xist deleted clones. While approximately 12-15% of genes were reactivated in △Xist-Pat-1 and △Xist-Pat-2 clones, △Xist-Pat-3 clone exhibited the reactivation of ∼40% of X-linked genes (Fig. 3A and 3C). When comparing the cohort of the reactivated genes among the three clones (△Xist-Pat-1, △Xist-Pat-2 and △Xist-Pat-3), we identified 20 X-linked genes that were reactivated across all three clones (Fig. 3C). Furthermore, many reactivated genes were unique to the △Xist-Pat-3 clone (Fig. 3C). Altogether, our analysis reveals that while many X-linked gene on the inactive X-chromosome can maintain their silent state in the absence of Xist, some are dependent on Xist.

**Figure 3:**
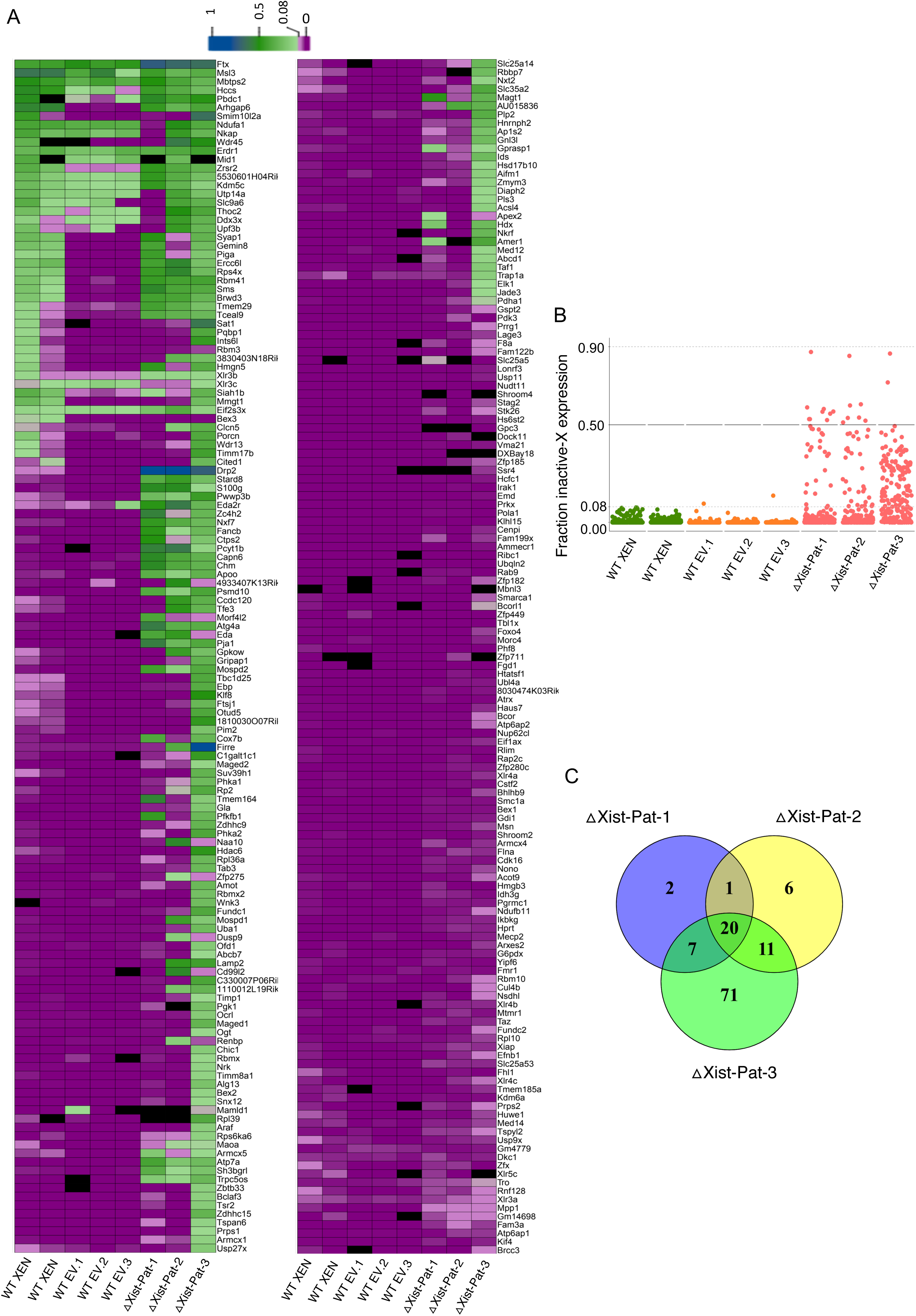
Paternal Xist deletion results in the partial reactivation of the inactive-X in XEN cells. (A) Heatmap showing the fraction of allelic expression of X-linked genes from the inactive-X in WT XEN, empty vector (WT EV.1, EV.2, and EV.3) and the heterozygous paternal Xist deleted clones (△Xist-Pat-1, △Xist-Pat-2 and △Xist-Pat-3). (B) The plot represents the fraction of inactive-X expression of reactivated X-linked genes. (C) Venn diagram representing the intersection of reactivated X-linked genes among three clones Xist-Pat-1, △Xist-Pat-2 and △Xist-Pat-3.

Similar to the imprinted X-inactivation, Xist plays significant role in orchestrating the initiation of random X-inactivation. However, the role of Xist in the maintenance of random X-chromosome inactivation remains poorly understood. In this study, we have used EpiSC as our model system to investigate the role of Xist in the maintenance of random X inactivation. EpiSC are the stem cells representing the post-implantation embryonic epiblast cells, which undergo random X-chromosome inactivation (Fig. 1A). The EpiSC line used for this study is a hybrid cell, and its two X-chromosomes are polymorphic in nature as they originated from two different mouse strains (X^Mus^: *Mus musculus* origin and X^Mol^: *Mus molossinus* origin). Moreover, this EpiSC line was isolated clonally from a single cell and therefore all cells harbored same inactivated X-chromosome (X^Mus^), which helped us to differentiate the expression of X-linked genes from active vs. inactive-X chromosome through allele specific analysis (Fig. 1A). To eliminate Xist expression in EpiSC cells, we employed the same CRISPR-Cas9 strategy as used for XEN cells. After considerable efforts, we were able to generate only one line of EpiSC (△Xist-Mus) with a heterozygous deletion of Xist from the inactive-X (X^Mus^) (Fig. 4A and 4B). The primary challenge we faced was the inefficient propagation of single cell clone, as EpiSC do not efficiently grow from a single cell. Next, to delineate the status of the inactive-X upon the loss of Xist, we sequenced whole transcriptome and performed allele-specific analysis. As expected, we observed that Xist was exclusively expressed from the X^Mus^ in WT EpiSC, thus validating further X^Mus^ is the inactive-X in this EpiSC line (Fig. 4C). Moreover, there was no expression of Xist in △Xist-Mus EpiSC, indicating complete loss of Xist expression in these cells (Fig. 4C). We then profiled the allelic expression fraction for X-linked genes originating from the inactive-X (X^Mus^). We found that many genes had almost no expression from the inactive-X (X^Mus^) in WT EpiSC, indicating robust X-chromosome inactivation (Fig. 4D). However, we found many genes escaped X-inactivation in WT EpiSC as indicated by their biallelic expression (Fig. 4D). Surprisingly, despite the loss of Xist in △Xist-Mus EpiSC, many genes maintained their silent state (Fig. 4D). Nevertheless, there was a subset of genes, which were reactivated upon loss of Xist in △Xist-Mus EpiSC (Fig. 4E). Taken together, we conclude that like XEN cells, the loss of Xist in EpiSC leads to the partial reactivation of the inactive-X chromosome.

**Figure 4:**
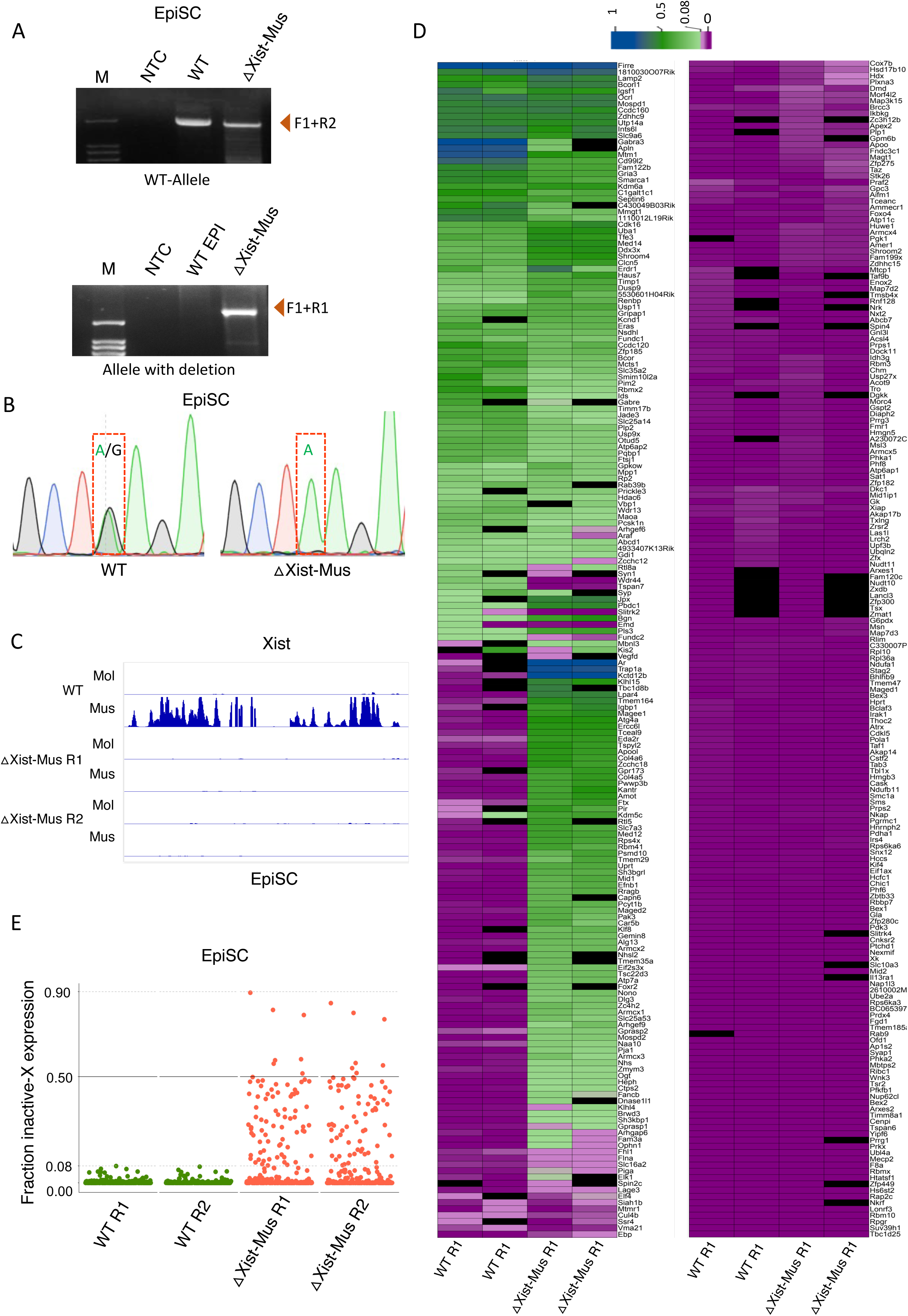
Ablation of Xist in EpiSC and profiling of the reactivation of X-linked genes. (A) PCR based genotyping of the *Xist* heterozygous deleted EpiSC line. WT allele was detected through F1+R2 primers, and deleted allele was detected through F1+R1 primers. M: marker, NTC: no template control. (B) Chromatogram showing the presence of inactive-X specific SNP in the deleted amplicon, indicating deletion of *Xist* from the inactive X. (C) Genome browser view of Xist locus showing no expression of Xist in △Xist-Mus EpiSC as profiled through allele-specific analysis of RNA-seq data. (D) Heatmap representing the fraction of allelic expression from the inactive-X chromosome in WT and △Xist-Mus EpiSC. (E) The plot represents the fraction of inactive X expression of the reactivated X-linked genes.

### Xist loss leads to the reactivation of same subset of X-linked genes in both XEN and EpiSC

Next, we performed a comparative analysis of the genes that were reactivated in △Xist XEN clones (△Xist-Pat-1, △Xist-Pat-2 and △Xist-Pat-3) with those in △Xist-Mus EpiSC. We considered only those genes, which expressed in △Xist XEN clones as well as △Xist-Mus EpiSC to avoid any potential confounding results due to the lack of expression in either group. Remarkably, we found that most of the reactivated genes in △Xist-Mus EpiSC (∼23 genes) overlapped with those in the △Xist XEN cell clones, only 5 genes, which were unique to EpiSC. This suggests that most of these genes are dependent on Xist to maintain their inactive state in both lineages (Fig. 5A). Subsequently, we explored if there is a connection between the kinetics of X-linked gene reactivation during the iPSC reprogramming and the susceptibility to the reactivation upon Xist loss in XEN and EpiSC. To investigate this, we compared our reactivated genes in △Xist XEN and △Xist EpiSC with the early, intermediate, and late reactivated genes reported in iPSCs reprogramming studies (32, 33) (Fig 5B and 5C). We observed that approximately 18 and 39 genes (for Talon and Janiszewski respectively), reactivated genes in △Xist XEN cells belonged to the early or intermediate classes of iPSC reprogramming. In contrast, about ∼12-13 genes belonged to the intermediate class. We also compared reactivated genes in △Xist XEN cells with the early, intermediate and late reactivated genes in germ cell reprogramming. We found that while around 25 genes were common with early reactivated genes, 32 genes belonged to late reactivated genes (Fig 5F). Together, it suggests that genes undergoing early, or intermediate reactivation are more prone to reactivation in XEN cells upon the loss of Xist (Fig 5B and 5C). Many other genes, which did not intersect could be due to lack of expression either in XEN and iPSC reprogramming cells (Fig 5B and 5C). On the other hand, several genes reactivated in △Xist EpiSC also belonged to the early/intermediate genes in iPSC reprogramming (6 to 15 genes in Talon and Janiszewski respectively) (Fig 5D and 5E). Six to nine genes belonged to the late reactivated class of iPSC reprogramming. However, when compared with the genes in germ cell reprogramming, only 5 genes intersected with the early/intermediate reactivated genes (Fig 5G). Six genes belonged to the late classes of germ cell reactivation (Fig 5G). Taken together, our analysis suggests that genes undergoing early to intermediate reactivation during cellular reprogramming are more susceptible to reactivation in both XEN and EpiSC upon thr loss of Xist.

**Figure 5:**
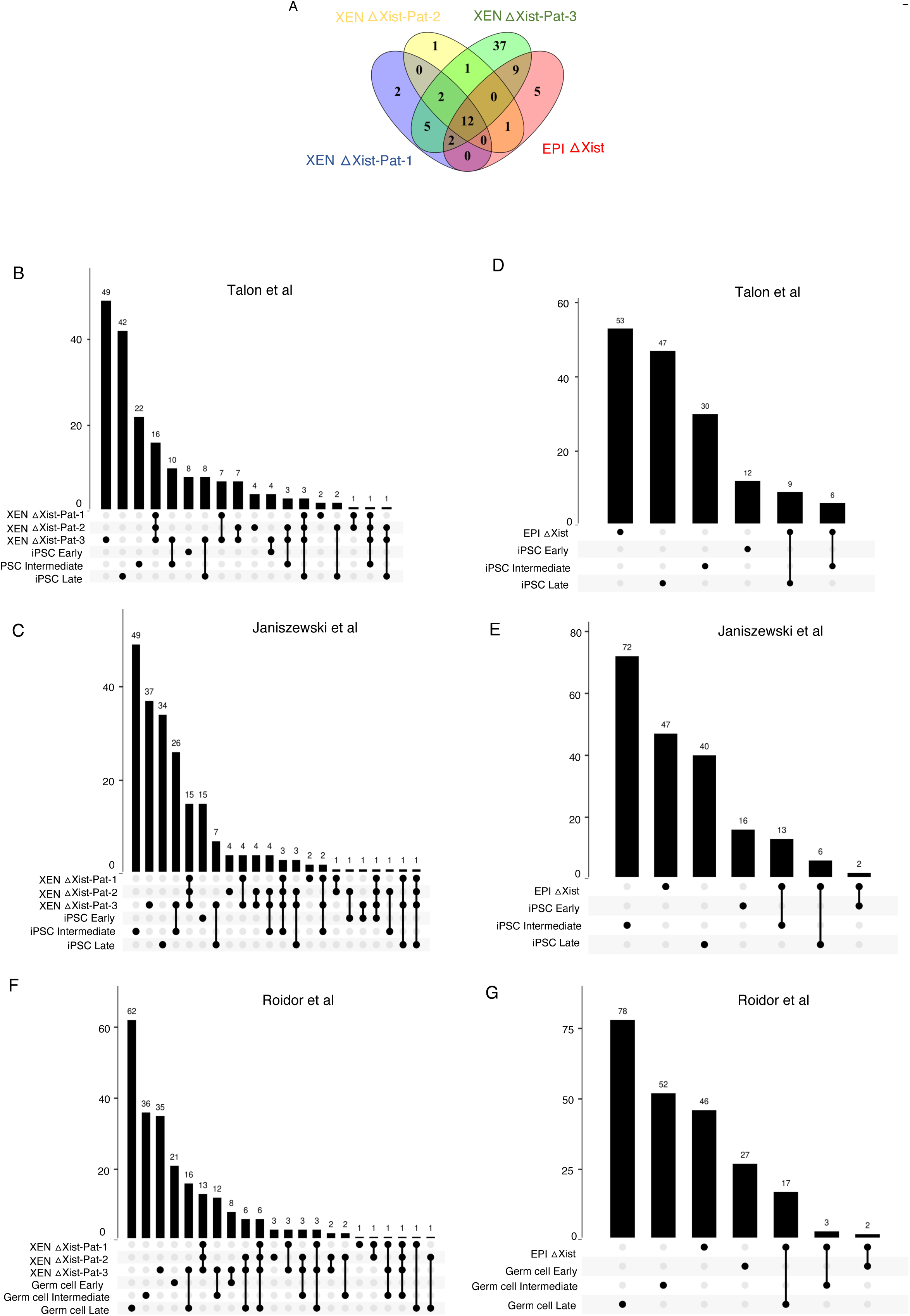
Corelating the X-linked gene reactivation to the reactivation kinetics during the reprogramming. (A)Venn diagram showing the intersection of the reactivated X-linked genes across Xist deleted XEN (△Xist-Pat-1, △Xist-Pat-2 △Xist-Pat-3 and △Xist-EpiSC. (B, C) Comparison of reactivated X-linked genes between △Xist XEN cells (△Xist-Pat-1, △Xist-Pat-2 △Xist-Pat-3) and iPSC reprogramming (early, intermediate and late). (D, E) Comparison of reactivated X-linked genes between △Xist EpiSC cells and iPSC reprogramming (early, intermediate and late). (F) Intersection plots representing the overlap between reactivated X-linked genes in △Xist XEN cells (△Xist-Pat-1, △Xist-Pat-2 △Xist-Pat-3) and reactivated genes of germ cell reprogramming (early, intermediate and late). (G) Intersection plots representing the overlap between reactivated X-linked genes in △Xist-EpiSC and reactivated genes of germ cell reprogramming (early, intermediate and late).

### Ablation of DNA methylation in Xist deleted XEN cells enhances the reactivation of the inactive-X

We hypothesized that CpG methylation at the promoter of X-linked genes could be a major contributing factor to the maintenance of the inactive state of X-linked genes that remained silent despite the loss of Xist in XEN cells. To test our hypothesis, we treated both WT and △Xist XEN clones (△Xist-Pat-1 and △Xist-Pat-2) with 5-Aza 2-deoxy-cytidine/Decitabine (5-AzaDC), an inhibitor of DNA methyl transferase DNMT1 for a period of around three days. Dot blot analysis of methylation level using a 5-methyl CpG (5-mc) antibody confirmed drastic reduction in DNA-methylation upon treatment with 5-AzaDC (Fig. 6A). Next, to profile the reactivation status of the inactive-X upon 5-AzaDC treatment, we conducted RNA-sequencing. As described above, we analyzed our RNA-seq data with allelic resolution to differentiate the expression between active vs. inactive-X and calculated the fraction of allelic expression originating from the inactive-X chromosome. Surprisingly, we found that 5-AzaDC treatment in WT XEN cells did not perturb X-linked gene silencing; only a few X-linked genes were reactivated. This suggests that CpG methylation is not essential for the maintenance of the XCI in XEN cells (Fig. 6B). However, 5-AzaDC treatment in △Xist XEN cells resulted in the reactivation of a subset of X-linked genes, which were not reactivated upon Xist loss. This implies that these genes depend on CpG methylation to maintain their inactive state in the absence of Xist (Fig. 6B). Nevertheless, many genes continued to maintain their silent state even after the loss of both Xist and DNA methylation, indicating the involvement of other unknown factors in the maintenance of XCI.

**Figure 6:**
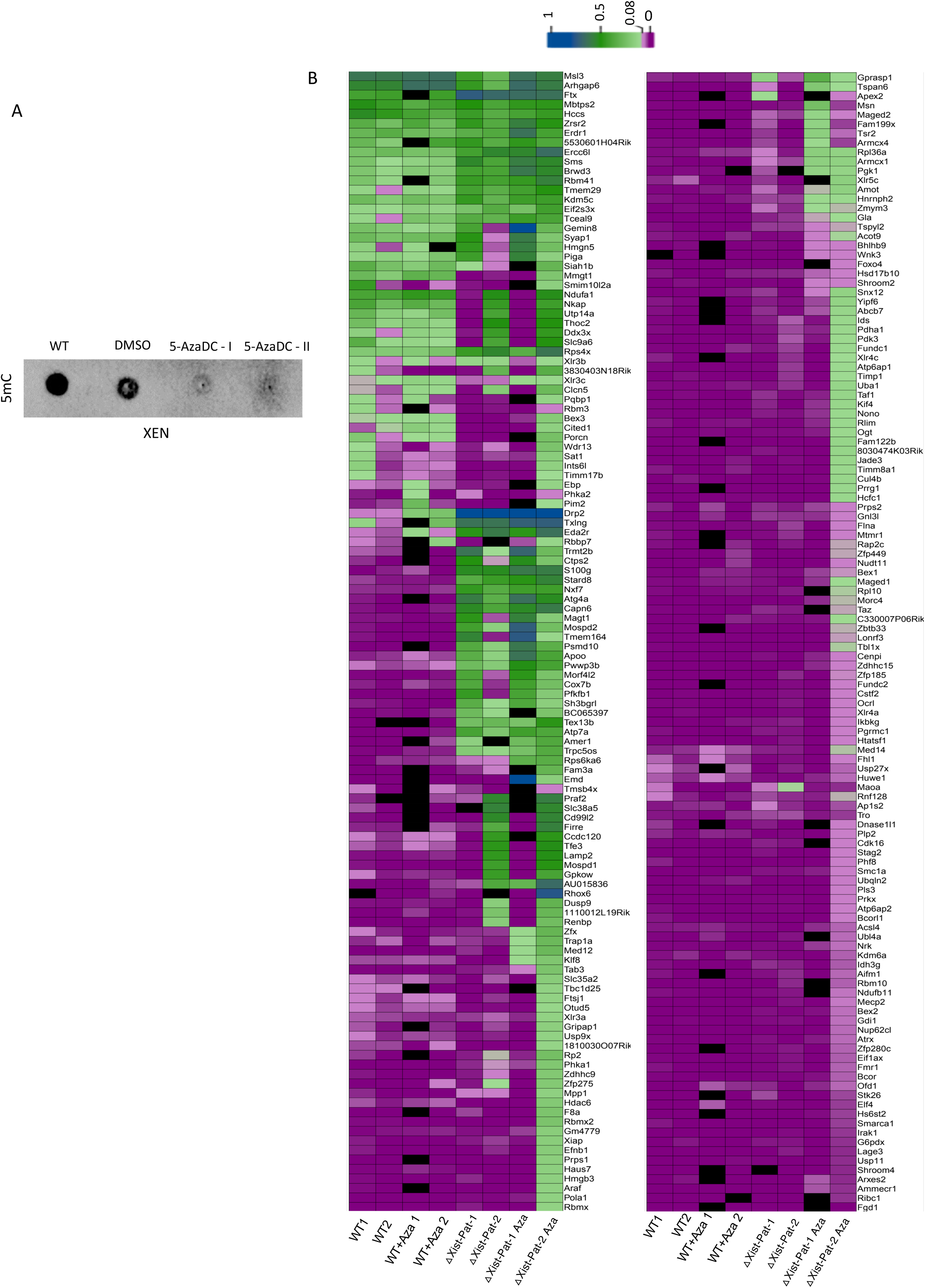
Ablation of DNA methylation and profiling of the reactivation of X-linked genes. (A) Representative dot blot showing the drastic reduction of DNA methylation upon treatment of Azacytidine in WT XEN cells. (B) Heatmap representing showing the reactivation of X-linked genes upon Azacytidine treatment in WT, △Xist-Pat-1, △Xist-Pat-2 cells. X-reactivation was profiled through calculating the fraction of allelic expression from the inactive-X chromosome through allele specific analysis of of RNA-seq data.

### Reactivation of X-linked genes upon Xist loss is associated with genomic and epigenomic features

Next, we investigated whether different genomic and epigenomic features could influence the potential for X-linked genes to be reactivated upon Xist loss. First, we tested if the location of the X-linked genes on the X-chromosome was linked to their reactivation potential. To explore this, we profiled the genomic distribution of the reactivated genes across the X-chromosome. We found that reactivated genes in △Xist-Pat-2 and △Xist-Pat-3 cells are located randomly across the X-chromosome, forming distinct small clusters (Fig. 7A). In contrast, △Xist-Pat-1 XEN and △Xist-EpiSC exhibited a different pattern (Fig. 7A). Reactivated genes in these cells were not evenly distributed but instead clustered in a specific region of the X-chromosome, near to the Xist locus (Fig. 7A). Moreover, clustering patterns were very similar between △Xist-Pat-1 XEN and △Xist-EpiSC cells (Fig. 7A). Notably, X-linked genes which were reactivated in all three △Xist-XEN clones also showed similar pattern (Fig. 7A). Taken together, we conclude that location of the genes on the X-chromosome often influences the reactivation potential of X-linked genes upon Xist loss.

**Figure 7:**
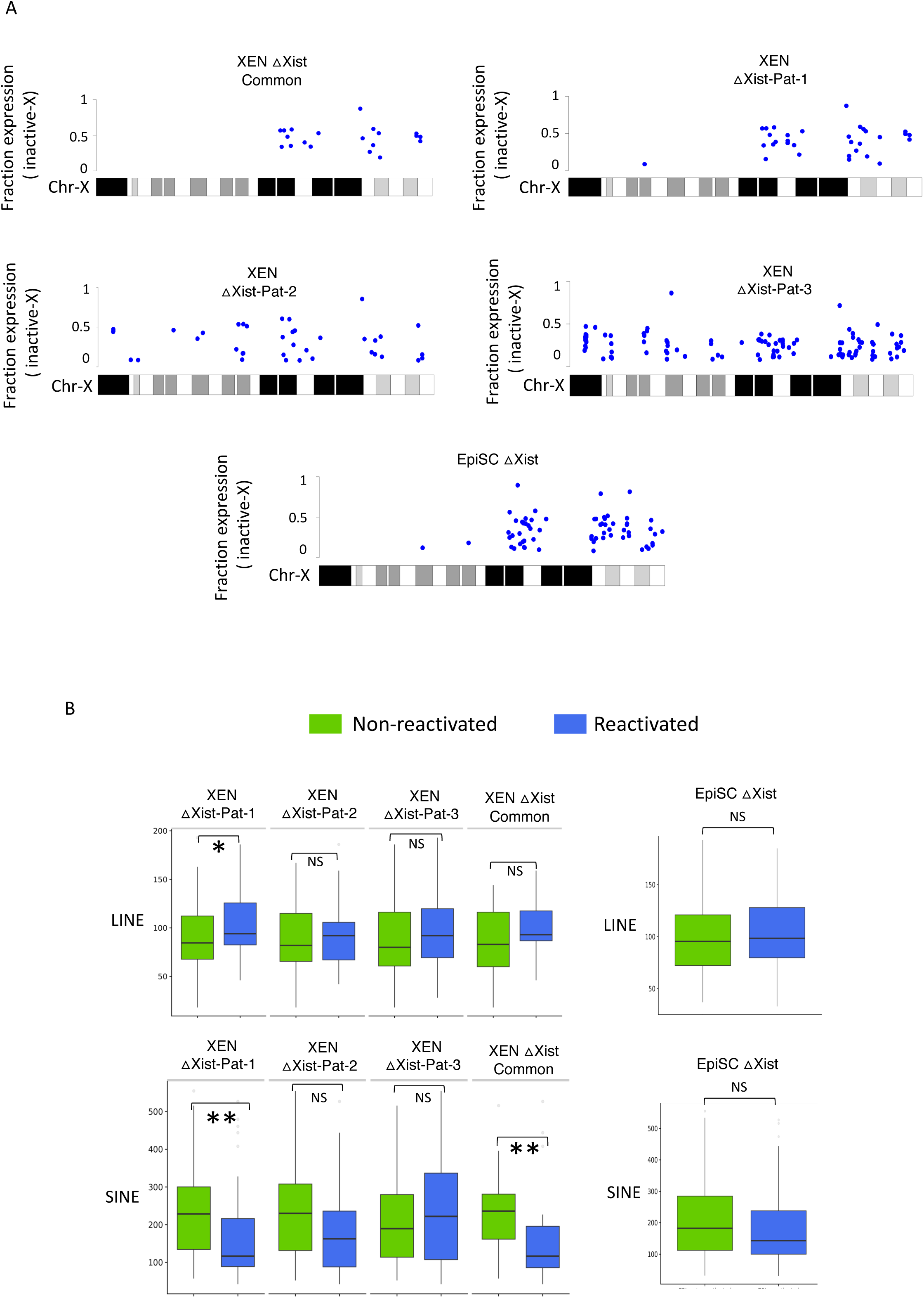
Profiling of genomic elements associated with the reactivated X-linked genes. (A) Schematic showing the location of the reactivated X-linked genes on the X-chromosome (B) Comparison of enrichment of the long interspaced nuclear elements (LINE, top) and short interspaced nuclear elements (SINE, bottom) between reactivated and non-reactivated genes.

Next, we compared the density of short-interspaced nuclear elements (SINEs) and long-interspaced nuclear elements (LINEs) in reactivated genes with non-reactivated X-linked genes (Fig 7B). Interestingly, reactivated genes tended to have higher LINEs compared to non-reactivated genes in △Xist-XEN and △Xist-EpiSC cells (Fig 7B). Conversely, reactivated genes showed less enrichment of SINE elements compared to non-reactivated genes (Fig 7B).

Subsequently, we delineated whether the chromatin state of the inactive-X genes played a role in determining the potential of reactivation of X-linked genes upon loss of Xist. To explore this, we performed ChIP-seq for different chromatin marks in WT XEN cells. We profiled the enrichment of different chromatin marks around the transcriptional start sites (TSSs) of the reactivated vs. non-reactivated X-linked gene loci on the inactive-X and active-X chromosome through allele-specific analysis of ChIP-seq data. Interestingly, we found that the reactivated genes showed lower enrichment of H3K9me3 compared to non-reactivated genes on the inactive-X chromosome, suggesting that genes with high H3K9me3 enrichment are less likely to be reactivated upon Xist loss (Fig. 8). In contrary, reactivated genes had slightly higher enrichment of H3K27me3 compared to non-reactivated genes at the inactive-X chromosome (Fig. 8). On the other hand, we found that active-chromatin marks such as H3K4me3 and H3K27ac were highly enriched at the homologous loci of reactivated genes compared to non-reactivated genes at the active-X chromosome. Other marks we profiled, such as H3K4me1, CTCF and Rad21 in XEN cells did not show any differential enrichment between reactivated vs. non-reactivated genes on either inactive-X or active-X chromosome (Fig. 8). Taken together, our analysis suggests that chromatin states play a pivotal role in determining the potential of the X-linked genes to be reactivated upon the loss of Xist.

**Figure 8:**
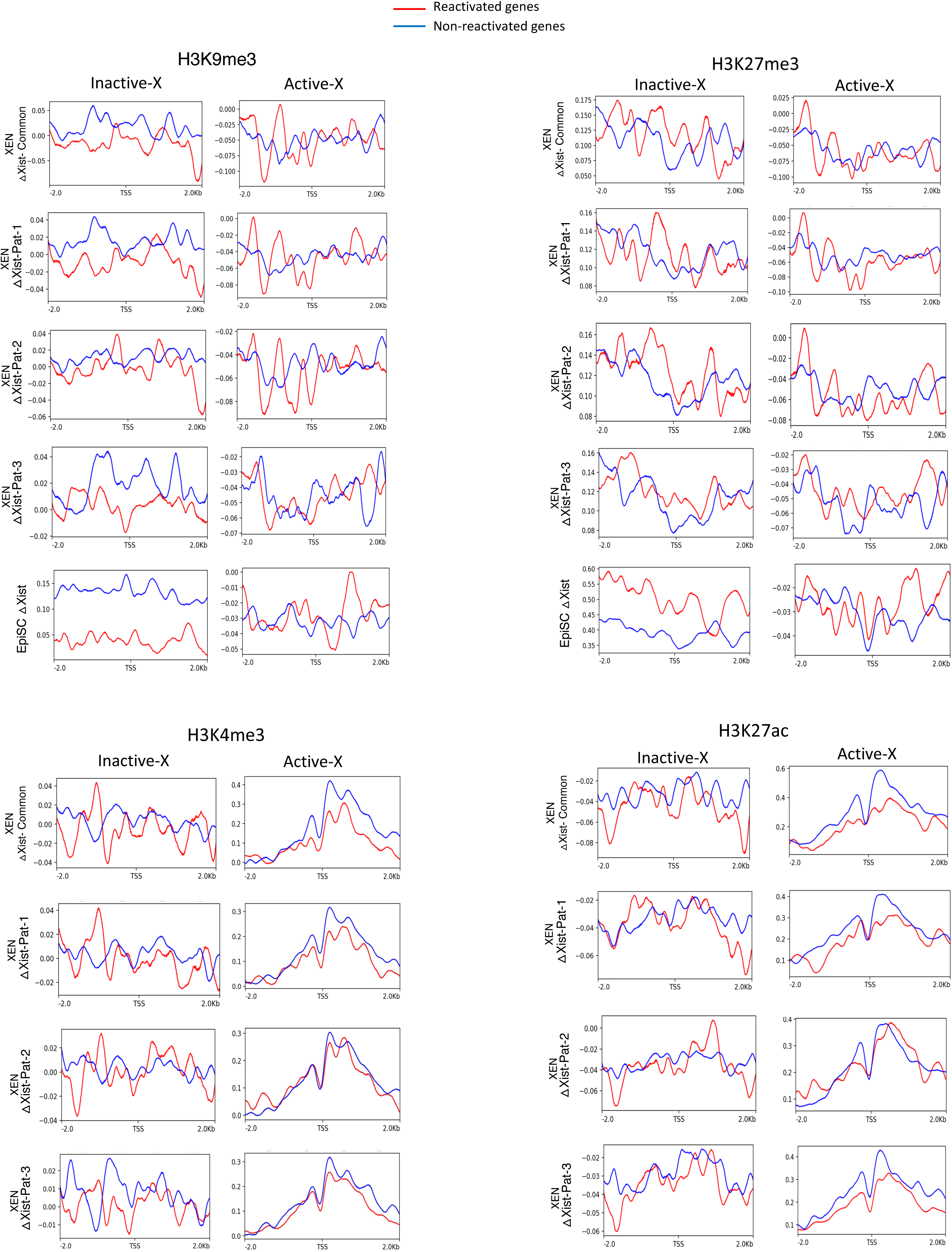
Profiling of chromatin states of reactivated vs. non-reactivated genes. Plots showing the enrichment of H3K9me3, H3K27me3, H3K4me3 and H3K27ac at TSS (+/-2Kb) of non-reactivated (blue) and reactivated genes (red) on the inactive and active X chromosome.

**Figure 9:**
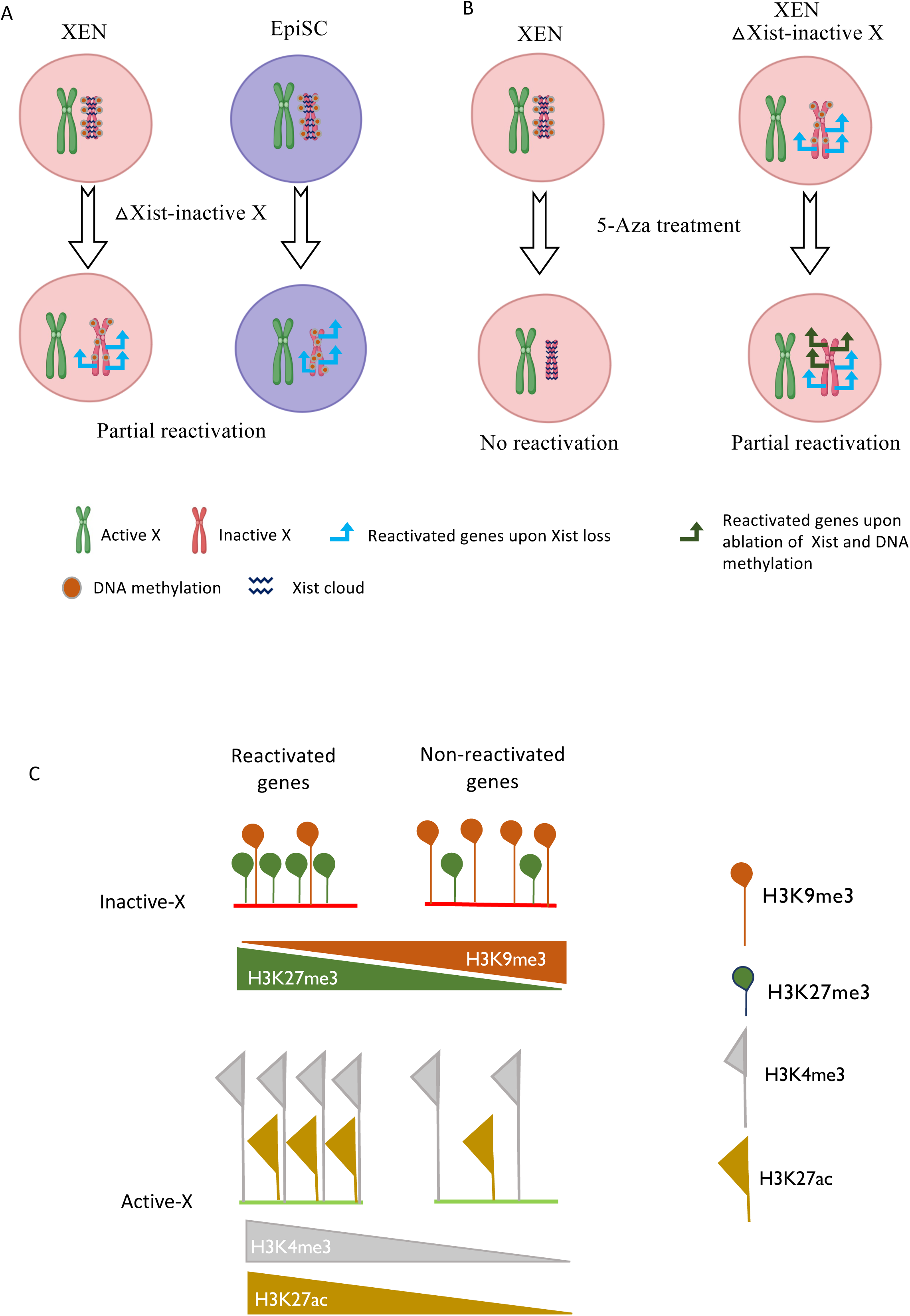
Working model. (A) Schematic showing that the Xist ablation in XEN and EpiSC leads to the partial reactivation of the inactive-X. Many genes reactivated in both cell types. (B) Ablation of DNA-methylation does not perturb X-inactivation maintenance in XEN cells. However, a subsets of X-linked genes gets reactivated when DNA methylation is ablated in Xist deficient XEN cells (C) Chromatin states are linked to the reactivation potential of X-linked genes upon Xist loss. Reactivated genes have high H3K27me3 and lesser H3K9me3 on the inactive-X. On the other hand, active-X counter parts of the reactivated genes are highly enriched with H3K4me3 and H3K27ac.

## Discussion

Emerging trends suggest that with aging, autoimmune diseases and cancer, often maintenance of XCI gets dysregulated, which often leads to the aberrant reactivation of many X-linked genes located on the inactive-X (34–38). However, the precise causes of this instability in the X-inactivation maintenance remains unknown. Therefore, it is crucial to investigate the factors and mechanisms responsible for maintaining the XCI. Here, we have explored into the factors and mechanisms that play important role in the maintenance of X-chromosome inactivation in early embryonic lineages: XEN and EpiSC. Our study highlights the integrated role of Xist, DNA-methylation and chromatin state play integrative roles in the maintenance of XCI.

We show that while most of the inactive-X genes can maintain their silennced state even in the absence of Xist RNA in Xist deleted XEN and EpiSC cells, a subset of genes undergo reactivation upon loss of Xist in these cells. This implies the involvement of other factors in the maintenance of gene silencing on the inactive-X (Fig. 3 and 4). Indeed, loss of Xist in other cell types such as hematopoietic stem cells, B-cells and mammary stem cells were also found to be associated with the partial reactivation of the inactive-X chromosome (28, 30, 39). Taken together, our study demonstrates that, like other cell types, early embryonic lineages are also partially rely on Xist RNA to maintain the inactivated state of X-linked genes and Xist contributes to the maintenance of XCI in a gene specific manner. Additionally, our study infers that like random X-inactivation, imprinted X-inactivation is also not entirely dependent on Xist to maintain the X-inactivation. Surprisingly, we found that the pool of reactivated genes in both XEN and EpiSC is mostly common, with only few exceptions. We conclude that these genes rely on Xist to maintain their inactive state irrespective of the lineages or type of XCI; i.e, imprinted or random. In contrary, earlier reports suggested that loss of Xist can have varying effects among different tissues or cell types (22, 29, 40). Moreover, we demonstrate that in XEN cells, the presence of Xist makes DNA methylation dispensable for the maintenance of XCI. However, in Xist deficient XEN cells, the loss of DNA methylation resulted in the reactivation of a subset of genes, which were not reactivated upon the loss of Xist (Fig. 6). This indicates that some genes depend on both Xist and DNA methylation to maintain their inactivated state.

On the other hand, we found that several genes reactivated upon Xist loss from both cell types of XEN and EpiSC, were belonged to early or intermediate reactivated genes during iPSC and germ cells reprogramming (Fig. 5). Based on this, we conclude that genes, which reactivate at the early or intermediate phases of cellular reprogramming are more susceptible to the reactivation in the absence of Xist. Furthermore, we found that reactivation potential of X-linked genes upon Xist loss is connected to their epigenomic and genomic features. We found that reactivated genes had low H3K9me3 enrichment and higher H3K27me3 enrichment compared to non-reactivated genes in both XEN and EpiSC (Fig.8). Previous studies in human induced pluripotent stem cells (iPSCs) have also shown that genes reactivated due to the X-erosion are highly enriched with H3K27me3 (21). Moreover, active-X counterpart of the reactivated genes were highly enriched with H3K4me3 and H3K27ac (Fig. 8). Moreover, we found that reactivated genes in △Xist-XEN and EpiSC tend to have higher LINEs element and lesser SINEs element. In contrary, a study in mouse neuronal stem cells showed that reactivated X-linked genes had fewer LINEs and higher SINEs compared to the non-reactivated X-linked genes in absence of Xist and CpG methylation (41).

### Materials and Method Data availability

Raw high throughput sequencing data (RNA-seq and ChIP-seq) generated in this study will be deposited to Gene Expression Omnibus (GEO) and accession will be provided soon after the publication. Upon request, data can be available by the corresponding author. The previously published dataset used for this study is available at GEO under the following accessions: GSE201189 (42).

### Cell culture

The XEN cells were cultured in the media composed of Dulbecco’s Modified Eagle medium (DMEM); (Himedia, #AL007A) supplemented with 20% fetal bovine serum (Invitrogen #10270106), 2mM L-glutamine (Invitrogen, #25030081), 100µM ß-mercaptoethanol (Sigma, #M3148), 100µM non-essential amino acids (NEAA; Invitrogen, #11140050) and penicillin-streptomycin (Gibco #15140122) in gelatin (Himedia # TCL059) coated plate at 37°C with 5% CO2. The cells were passaged using 0.05% trypsin EDTA (Himedia #TCL033).

EpiSC were grown in Matrigel (Corning, #354277) coated plate in knockout-DMEM (Gibco, #10829018) supplemented with 20% knockout fetal bovine serum (Invitrogen #10828028), 2mM L-glutamine (Invitrogen, #25030081), 100µM ß-mercaptoethanol (Sigma #M3148), 100µM non-essential amino acids (Invitrogen, #11140050), 10ng/ml fibroblast growth factor 2 (Prospec #CYT 386) and penicillin-streptomycin (Gibco #15140122). The cells were passaged with 0.05% trypsin EDTA (Himedia #TCL033).

### Generation of Xist Knockout in XEN and EpiSC by CRISPR/Cas9

The dual sgRNAs (SgRNA1: AATCCGGAGTATGAGCTGAGAGG SgRNA2: TGCTGCGAAGGAATT TACAGTGG) targeting the Xist at the promoter and intron1 were designed using chop-chop (https://chopchop.cbu.uib.no/ <u>red font nucleotides were for spCas9</u>). The sgRNAs were cloned in to the pSpCas9(BB)-2A-Puro V2.0 (Px459; Add gene, #62988) plasmid. The XEN/EpiSC cells were transfected with Px459-sgRNA plasmids using the lipofectamine 2000 (Thermo Scientific, #11668019) and were selected with 3µg/ml of puromycin (Takara, #631305). After selection, the transfected cells were cultured in the fresh media without puromycin unless the colonies were observed. Next, we plated putative Xist deleted XEN cells as a single-cell into a 96-well plate using fluorescent associated cell sorter (FACS). The putative Xist deleted EpiSC were plated as single cell in 96 well plate using manual method and grown in Cellartis DEF CS 500 culture system (Takara, #Y30012). Next, single cell clones were grown, and DNA was isolated using Direct PCR tail lysis buffer (Viagen, V102T). Finally, clones were screened PCR and Sanger sequencing using the screening primers (F1: CTGGGACTAGAGACATACCT, R1: ACTTCCACTTCCCACCACTTA, R2: TAACACCTCAGGGACAGGAA).

### 5-Aza deoxycytidine treatment

To deplete DNA-methylation, WT XEN and △Xist XEN cells were treated with 5-Aza 2 - deoxy-cytidine/Decitabine (5AzaDC, Sigma, #A3656), an inhibitor of DNA methyl transferase DNMT1. In brief, appropriate stock solution of 5AzaDC was prepared in dimethyl sulfoxide (DMSO) (Sigma #A2438). Stock solution was further diluted in culture media to treat XEN cells with the final concentration of 20µM. Cells were treated for 70 hours by changing the media with fresh drug for every 24 hours. After 70 hours, the cells were harvested for the DNA and RNA isolation.

### DNA isolation and Dot-blot

Genomic DNA was isolated from WT and 5AzaDC treated XEN cells using blood and cell culture DNA mini kit (Qiagen #Q13323) following manufacturers protocol. In brief, approximately 5 × 10^6^ cells were harvested, washed with 1X Dulbecco’s phosphate buffer saline (1X DPBS, Hi-media, TL1006), and then cells were lysed with ice-cold lysis buffer. Next, nuclei were pelleted down through centrifugation at 1300 X g at 4° C for 10 min. Nuclei was dissolved with G2 buffer along with proteinase K for 30 min at 50° C and DNA was extracted through the column with the elution buffer. The eluted DNA was precipitated with the iso-propanol (Qualigens #13825), washed with ice-cold 70% ethanol (Hayman #F205220) followed by vacuum dried. Finally, DNA was resuspended in ultrapure water through incubating at 55°C for 1 hour while shaking at 300 rpm.

For the dot blot assay, the genomic DNA was fragmented (200 – 400 bps) through sonication with Biorupter pico (Diagenode) for 5 cycles 20 sec on and 70 sec off. Next, DNA was heat denatured at 95°C for 10 min. Denatured DNA was spotted on the nylon membrane (GE Health care # RPN303B) with equal amount of DNA (20ng) and air dried. After UV-crosslinking, membrane was blocked with the 5% bovine serum albumin (BSA). Next, membrane was incubated with 1°Ab against 5-methyl CpG (CST # 28692S,) at 1:2000 dilution at 4°C (cold room) for overnight. Next day, the membrane was washed with 1X TBS for 3 times with each 5 minutes incubation at room temperature. Then membrane was incubated with the HRP conjugated 2°Ab (Invitrogen # 31460) at 1:5000 at room temperature for 2 hours. After washing the membrane with 1X TBS – for 3 times with each 5 minutes incubation at room temperature, the blot was developed using ECL (Invitrogen # 35480).

### RNA-FISH

RNA-FISH was performed as described previously (43). Briefly, double-stranded RNA-FISH probes were generated through labelling BAC DNA (Xist, Atrx, Rlim and Pgk1) using Bio Prime™ labeling kit (Invitrogen, # 18094-011) following the manufactures protocol. Probes were labelled either with Cy3-dUTP (Enzo Life Sciences, #ENZ471 42501) or Cy5-dUTP. After labelling reaction completed, probes were purified through ProbeQuant G-50 Micro columns (Cytiva,472 #28903408). Labelled probes were then precipitated using 0.3M sodium acetate (Sigma, #71196) along with 300 µg of Yeast tRNA (Invitrogen, #15401011), 150 µg of sheared Salmon sperm DNA (Invitrogen, #15632-011) and absolute ethanol (Hayman, #F205220) through centrifuging at 13,000 rpm for 20 mins at 4°C. Following the precipitation, the pellet was washed with 70% and then 100% ethanol and thereafter vacuum-dried. Next, probes were resuspended in deionized formamide (VWR Life Sciences, #0606) and then denatured at 95°C for 10 mins followed by immediate snap-freezing on ice. A hybridization solution was prepared with 20% Dextran sulphate (SRL, #76203), 2X SSC (SRL, #12590) and was added to the denatured probes.

Briefly, cells were cultured and grown on gelatin coated coverslips. After reaching to appropriate confluency, cells were permeabilized with ice-cold Cytoskeletal (CSK) buffer (300 mM Sucrose, 100mM NaCl, 3mM MgCl2 and 10mM PIPES buffer, pH 6.8) and Triton X-100 Following the permeabilization, cells were fixed using 3% paraformaldehyde (Electron Microscopy Sciences, #15710) for 10 min. Before hybridization, cells were dehydrated through ethanol series 70%, 85%, 95% and 100% incubating each for 2 min, individually. Subsequently, coverslip was allowed to air dry for 15 min at room temperature. Next, hybridization with the labelled probe was set up in a humid chamber containing a solution of 50% Formamide (MP Biomedicals, #193995) and 2X SSC at 37°C for overnight. Next day, coverslips were washed with 2X SSC/50% formamide three times, followed by three washes with 2X SSC, and finally two washes with 1X SSC for 7 mins each at 37°C. DAPI (Invitrogen, #D1306) was added during the 2X SSC wash. After completion of the washing, coverslips were mounted using Vectashield (Vector Labs, #H1000) and sealed with nail polish. before analysis under microscope.

### Microscopy

Image acquisition for RNA-FISH was performed using Olympus IX73 Inverted microscope (Olympus LS) build with a camera using 100X objective and using cellSens [Ver.2.1] Life Science imaging software (Olympus LS).

### RNA sequencing and analysis

For RNA-sequencing, cells were harvested in Trizol (Life technologies #15596-026) and RNA was isolated according to the manufacture’s method. Before library preparation, quality of RNA was checked through gel electrophoresis as well as through Bioanalyzer (Agilent technologies). RNA with RNA integrity number (RIN) above 7 was considered for the library preparation. NEBNext Ultra/Tru-seq RNA library preparation kit compatible with Illumina sequencing were used for the library preparation. Library was sequenced through Illumina platform following paired end (2 × 150 bp) chemistry. For analysis of RNA-sequencing data, reads quality were checked through FastQC and adaptor trimming was performed using Trim Galore (v0.6.6). Any ribosomal RNA content was removed through RiboDetector (v0.2.7) (44). Reads were mapped to the mouse reference genome GRCm38 (mm10) using STAR (2.7.9a) and counting was carried out using featureCounts (2.0.1) tool (45, 46).

### Allele specific RNA-seq analysis

To profile the allelic expression of X-linked genes, we analysed RNA-seq data with allelic resolution. Allele specific RNA-seq analysis was carried out as described earlier (31, 47, 48). In brief, we leveraged strain specific SNPs (Mus musculus musculus (Mus) and Mus musculus molossinus (Mol)) to perform allele-specific analysis. Strain-specific SNPs were obtained from the Mouse Genomes Project (https://www.sanger.ac.uk/data/mouse-genomes-project/). Two separate strain-specific in silico reference genomes were constructed by placing strain-specific SNPs into the mouse GRCm38 (mm10) reference genome using variant calling file (VCF) tools. Next, reads were mapped to these in silico constructed strain specific reference genomes using STAR. Following the allele-specific mapping pf the reads, SNP-wise reads were counted. Only those SNPs were considered for our analysis, which had at least 10 reads per SNP-site (Mus + Mol). For determining of gene wise allelic read counts, we filtered those genes, which had at least 2 SNPs. Allelic reads of a gene were calculated taking average of SNP-wise reads. Fraction allelic expression was estimated through following formula: Allele Mus or Allele Mol/Allele Mus + Allele Mol.

### ChIP-sequencing

ChIP was performed using the SimpleChIP Enzymatic Chromatin IP kit (Cell Signaling Technology, #9003) as described earlier. In brief, appropriate number of cells were fixed with formaldehyde and quenched with 125mM glycine. Next, cells were scraped with cell scraper and lysed with the lysis buffer provided in the kit. After that, chromatin was fragmented using 0.5 μl micrococcal nuclease per IP followed by sonication with Biorupter pico (Diagenode) for 20 cycles 30 sec on/off. Immunoprecipitation was done using ∼1ug antibody of CTCF (Diagenode, #C15410210), Rad21 (Abcam, #ab992), H3K4me1 (Diagenode, #C15410194), H3K27ac (Diagenode, #C15410196), H3K4me3 (Diagenode), H3K9me3 (Diagenode) and H3K27me3 (Millipore). In parallel, immunoprecipitation was done for isotype control IgG (provided in the kit). 2% input sample was kept aside before the immunoprecipitation, and which was used for the normalization of the ChIP signal. After reverse crosslinking, the immunoprecipitated DNA was purified and quantified using Qubit fluorometer (Thermo Fisher Scientific, USA). Next, ChIP-seq library was prepared using NEXTflex Rapid DNA sequencing Bundle kit (BIOO Scientific, Inc. U.S.A.). Library was sequence in Illumina Hiseq platform using paired end (150 × 2) chemistry.

### Allele-specific ChIP-seq analysis

Allele-specific analysis of ChIP-seq data was performed as described previously (49). First, using SNPsplit genome preparation, a N-masked in-silico genome was created (50). Followed by, ChIP-seq reads were mapped to the N-masked genome using Bowtie2 (51). Next, strain specific BAM files were generated by allelic segregation of reads using SNPsplit (0.4.0). Enrichment ratio log2 (ChIP/input) was calculated using bamCompare from deeptools (v.3.5.0) with BPM normalization (52).

## Acknowledgement

We thank Prof. Sundeep Kalantry, University of Michigan for gifting BACs and cell lines used in this study. We acknowledge the FACS facility of department of developmental biology and genetics at Indian institute of scienece, Bangalore, India. This study is supported by the Department of Biotechnology (DBT), Govt. of India grant (BT/PR30399/BRB/10/1746/2018), DST-SERB (CRG/2019/003067), DBT-Ramalingaswamy fellowship (BT/RLF/Re-entry/05/2016) and Infosys Young Investigator grant award to SG. SM acknowledge the fellowship support from University Grants Commision (UGC). RB acknowledge Council of Scientific and Industrial Research (CSIR), India for the fellowship. LSB acknowledge DST-Women scientist award.

## Authors Contribution

SG supervised and acquired the funding for the study. SG and MA conceptualized the study. MA, SM, LSB, RB, performed experiments. HCN performed bioinformatic analysis. MA wrote the first draft of the manuscript. SG and MA edited and proofread the manuscript. The final manuscript was approved by all the authors.

## References

1. Lyon, M.F. (1961) Gene action in the X-chromosome of the mouse (mus musculus L.). Nature, 190, 372–373.

2. Maclary, E., Buttigieg, E., Hinten, M., Gayen, S., Harris, C., Sarkar, M.K., Purushothaman, S. and Kalantry, S. (2014) Differentiation-dependent requirement of Tsix long non-coding RNA in imprinted X-chromosome inactivation. Nat. Commun., 5.

3. Maclary, E., Hinten, M., Harris, C., Sethuraman, S., Gayen, S. and Kalantry, S. (2017) PRC2 represses transcribed genes on the imprinted inactive X chromosome in mice. Genome Biol., 18.

4. Harris, C., Cloutier, M., Trotter, M., Hinten, M., Gayen, S., Du, Z., Xie, W. and Kalantry, S. (2019) Conversion of random x-inactivation to imprinted x-inactivation by maternal prc2. Elife, 8.

5. Gayen, S., Maclary, E., Buttigieg, E., Hinten, M. and Kalantry, S. (2015) A Primary Role for the Tsix lncRNA in Maintaining Random X-Chromosome Inactivation. Cell Rep., 11, 1251–1265.

6. Monkhorst, K., Jonkers, I., Rentmeester, E., Grosveld, F. and Gribnau, J. (2008) X Inactivation Counting and Choice Is a Stochastic Process: Evidence for Involvement of an X-Linked Activator. Cell, 132, 410–421.

7. Gayen, S., Maclary, E., Hinten, M. and Kalantry, S. (2016) Sex-specific silencing of X-linked genes by Xist RNA. Proc. Natl. Acad. Sci., 113, E309–E318.

8. Samanta, M.K., Gayen, S., Harris, C., Maclary, E., Murata-Nakamura, Y., Malcore, R.M., Porter, R.S., Garay, P.M., Vallianatos, C.N., Samollow, P.B., et al. (2022) Activation of Xist by an evolutionarily conserved function of KDM5C demethylase. Nat. Commun., 13, 2602.

9. Chu, C., Zhang, Q.C., Da Rocha, S.T., Flynn, R.A., Bharadwaj, M., Calabrese, J.M., Magnuson, T., Heard, E. and Chang, H.Y. (2015) Systematic discovery of Xist RNA binding proteins. Cell, 161, 404–416.

10. Minajigi, A., Froberg, J.E., Wei, C., Sunwoo, H., Kesner, B., Colognori, D., Lessing, D., Payer, B., Boukhali, M., Haas, W., et al. (2015) A comprehensive Xist interactome reveals cohesin repulsion and an RNA-directed chromosome conformation. Science (80-.)., 349, 1DUIMMY.

11. Weber, M., Hellmann, I., Stadler, M.B., Ramos, L., Pääbo, S., Rebhan, M. and Schübeler, D. (2007) Distribution, silencing potential and evolutionary impact of promoter DNA methylation in the human genome. Nat. Genet., 39, 457–466.

12. Marahrens, Y., Panning, B., Dausman, J., Strauss, W. and Jaenisch, R. (1997) Xist-deficient mice are defective in dosage compensation but not spermatogenesis. Genes Dev., 11, 156–166.

13. Yang, L., Kirby, J.E., Sunwoo, H. and Lee, J.T. (2016) Female mice lacking Xist RNA show partial dosage compensation and survive to term. Genes Dev., 30, 1747–1760.

14. Marahrens, Y., Loring, J. and Jaenisch, R. (1998) Role of the Xist Gene in X Chromosome Choosing. Cell, 92, 657–664.

15. Brown, C.J. and Willard, H.F. (1994) The human X-inactivation centre is not required for maintenance of X-chromosome inactivation. Nature, 368, 154–156.

16. Csankovszki, G., Panning, B., Bates, B., Pehrson, J.R. and Jaenisch, R. (1999) Conditional deletion of Xist disrupts histone macroH2A localization but not maintenance of X inactivation [1]. Nat. Genet., 22, 323–324.

17. Csankovszki, G., Nagy, A. and Jaenisch, R. (2001) Synergism of Xist RNA, DNA methylation, and histone hypoacetylation in maintaining X chromosome inactivation. J. Cell Biol., 153, 773–783.

18. Wutz, A. and Jaenisch, R. (2000) A shift from reversible to irreversible X inactivation is triggered during ES cell differentiation. Mol. Cell, 5, 695–705.

19. Lee, H.J., Gopalappa, R., Sunwoo, H., Choi, S.-W., Ramakrishna, S., Lee, J.T., Kim, H.H. and Nam, J.-W. (2019) En bloc and segmental deletions of human XIST reveal X chromosome inactivation-involving RNA elements. Nucleic Acids Res., 10.1093/nar/gkz109.

20. Lv, Q., Yuan, L., Song, Y., Sui, T., Li, Z. and Lai, L. (2016) D-repeat in the XIST gene is required for X chromosome inactivation. RNA Biol., 13, 172–176.

21. Vallot, C., Ouimette, J.F., Makhlouf, M., Féraud, O., Pontis, J., Côme, J., Martinat, C., Bennaceur-Griscelli, A., Lalande, M. and Rougeulle, C. (2015) Erosion of X chromosome inactivation in human pluripotent cells initiates with XACT coating and depends on a specific heterochromatin landscape. Cell Stem Cell, 16, 533–546.

22. Yildirim, E., Kirby, J.E., Brown, D.E., Mercier, F.E., Sadreyev, R.I., Scadden, D.T. and Lee, J.T. (2013) Xist RNA is a potent suppressor of hematologic cancer in mice. Cell, 152, 727–742.

23. Bhatnagar, S., Zhu, X., Ou, J., Lin, L., Chamberlain, L., Zhu, L.J., Wajapeyee, N. and Green, M.R. (2014) Genetic and pharmacological reactivation of the mammalian inactive X chromosome. Proc. Natl. Acad. Sci., 111, 12591–12598.

24. Zhang, L.F., Huynh, K.D. and Lee, J.T. (2007) Perinucleolar Targeting of the Inactive X during S Phase: Evidence for a Role in the Maintenance of Silencing. Cell, 129, 693– 706.

25. Carrette, L.L.G., Wang, C.Y., Wei, C., Press, W., Ma, W., Kelleher, R.J. and Lee, J.T. (2018) A mixed modality approach towards Xi reactivation for Rett syndrome and other X-linked disorders. Proc. Natl. Acad. Sci. U. S. A., 115, E668–E675.

26. Anguera, M.C., Sadreyev, R., Zhang, Z., Szanto, A., Payer, B., Sheridan, S.D., Kwok, S., Haggarty, S.J., Sur, M., Alvarez, J., et al. (2012) Molecular Signatures of Human Induced Pluripotent Stem Cells Highlight Sex Differences and Cancer Genes. Cell Stem Cell, 11, 75–90.

27. Yang, T., Ou, J. and Yildirim, E. (2022) Xist exerts gene-specific silencing during XCI maintenance and impacts lineage-specific cell differentiation and proliferation during hematopoiesis. Nat. Commun., 13.

28. Richart, L., Picod-Chedotel, M.L., Wassef, M., Macario, M., Aflaki, S., Salvador, M.A., Héry, T., Dauphin, A., Wicinski, J., Chevrier, V., et al. (2022) XIST loss impairs mammary stem cell differentiation and increases tumorigenicity through Mediator hyperactivation. Cell, 185, 2164–2183.e25.

29. Adrianse, R.L., Smith, K., Gatbonton-Schwager, T., Sripathy, S.P., Lao, U., Foss, E.J., Boers, R.G., Boers, J.B., Gribnau, J. and Bedalov, A. (2018) Perturbed maintenance of transcriptional repression on the inactive X-chromosome in the mouse brain after Xist deletion. Epigenetics and Chromatin, 11.

30. Yu, B., Qi, Y., Li, R., Shi, Q., Satpathy, A.T. and Chang, H.Y. (2021) B cell-specific XIST complex enforces X-inactivation and restrains atypical B cells. Cell, 184, 1790–1803.e17.

31. Naik, H., Chandel, D., Majumdar, S., Arava, M., Baro, R., Bv, H., Hari, K., Parichitran, Avinchal, Jolly, M., et al. (2023) Lineage-specific dynamics of erasure of X-upregulation during inactive-X reactivation. bioRxiv, 10.1101/2020.12.23.424181.

32. Talon, I., Janiszewski, A., Theeuwes, B., Lefevre, T., Song, J., Bervoets, G., Vanheer, L., De Geest, N., Poovathingal, S., Allsop, R., et al. (2021) Enhanced chromatin accessibility contributes to X chromosome dosage compensation in mammals. Genome Biol., 22.

33. Janiszewski, A., Talon, I., Chappell, J., Collombet, S., Song, J., De Geest, N., Kit To, S., Bervoets, G., Marin-Bejar, O., Provenzano, C., et al. (2019) Dynamic reversal of random X-Chromosome inactivation during iPSC reprogramming. Genome Res., 29, 1659–1672.

34. Chaligné, R., Popova, T., Mendoza-Parra, M.A., Saleem, M.A.M., Gentien, D., Ban, K., Piolot, T., Leroy, O., Mariani, O., Gronemeyer, H., et al. (2015) The inactive X chromosome is epigenetically unstable and transcriptionally labile in breast cancer. Genome Res., 25, 488–503.

35. Kang, J., Lee, H.J., Kim, J., Lee, J.J. and Maeng, L.S. (2015) Dysregulation of X chromosome inactivation in high grade ovarian serous adenocarcinoma. PLoS One, 10.

36. Pageau, G.J., Hall, L.L., Ganesan, S., Livingston, D.M. and Lawrence, J.B. (2007) The disappearing Barr body in breast and ovarian cancers. Nat. Rev. Cancer, 7, 628–633.

37. Basu, A., Syrett, C.M., Anguera, M.C., Kramer, M.C., Wang, J. and Atchison, M.L. (2016) Unusual maintenance of X chromosome inactivation predisposes female lymphocytes for increased expression from the inactive X. Proc. Natl. Acad. Sci., 113, E2029–E2038.

38. Wareham, K.A., Lyon, M.F., Glenister, P.H. and Williams, E.D. (1987) Age related reactivation of an X-linked gene. Nature, 10.1038/327725a0.

39. Yang, T., Ou, J. and Yildirim, E. (2022) Xist exerts gene-specific silencing during XCI maintenance and impacts lineage-specific cell differentiation and proliferation during hematopoiesis. Nat. Commun., 13.

40. Yang, L., Kirby, J.E., Sunwoo, H. and Lee, J.T. (2016) Female mice lacking Xist RNA show partial dosage compensation and survive to term. Genes Dev., 30, 1747–1760.

41. Mira-Bontenbal, H., Tan, B., Gontan, C., Goossens, S., Boers, R.G., Boers, J.B., Dupont, C., van Royen, M.E., IJcken, W.F.J., French, P., et al. (2022) Genetic and epigenetic determinants of reactivation of Mecp2 and the inactive X chromosome in neural stem cells. Stem Cell Reports, 17, 693–706.

42. Ichihara, S., Nagao, K., Sakaguchi, T., Obuse, C. and Sado, T. (2022) SmcHD1 underlies the formation of H3K9me3 blocks on the inactive X chromosome in mice. Development, 149.

43. Majumdar, S., Bammidi, L., Naik, H., Avinchal, Baro, R., Kalita, A., Sundarraj, N., Bariha, G., Notani, D. and Gayen, S. (2023) Deletion of Xist upstream sequences alters TAD interactions and leads to defects in Xist coating and expression. bioRxiv, 10.1101/2023.08.14.553118.

44. Deng, Z.L., Münch, P.C., Mreches, R. and McHardy, A.C. (2022) Rapid and accurate identification of ribosomal RNA sequences via deep learning. Nucleic Acids Res., 50, e60–e60.

45. Dobin, A., Davis, C.A., Schlesinger, F., Drenkow, J., Zaleski, C., Jha, S., Batut, P., Chaisson, M. and Gingeras, T.R. (2013) STAR: Ultrafast universal RNA-seq aligner. Bioinformatics, 29, 15–21.

46. Liao, Y., Smyth, G.K. and Shi, W. (2014) featureCounts: an efficient general purpose program for assigning sequence reads to genomic features. Bioinformatics, 30, 923–930.

47. Naik, H.C., Hari, K., Chandel, D., Jolly, M.K. and Gayen, S. (2022) Single-cell analysis reveals X upregulation is not global in pre-gastrulation embryos. iScience, 25.

48. Naik, H.C., Hari, K., Chandel, D., Mandal, S., Jolly, M.K. and Gayen, S. (2021) Semicoordinated allelic-bursting shape dynamic random monoallelic expression in pregastrulation embryos. iScience, 24, 102954.

49. Parichitran, A., Naik, H., Naskar, A., Bammidi, L. and Gayen, S. (2023) Chromatin states contribute to coordinated allelic transcriptional bursting 1 to drive iPSC reprogramming 2 3 Parichitran A. bioRxiv, 10.1101/2023.07.13.548864.

50. F, K. and SR, A. (2016) SNPsplit: Allele-specific splitting of alignments between genomes with known SNP genotypes. F1000Research, 5.

51. B, L. and SL, S. (2012) Fast gapped-read alignment with Bowtie 2. Nat. Methods, 9, 357–359.

52. Ramírez, F., Ryan, D.P., Grüning, B., Bhardwaj, V., Kilpert, F., Richter, A.S., Heyne, S., Dündar, F. and Manke, T. (2016) deepTools2: a next generation web server for deep-sequencing data analysis. Nucleic Acids Res., 44, W160–W165.

