## Supplemental figures for "Gene-specific reactivation of X-linked genes upon Xist loss is linked to the chromatin states in extraembryonic endoderm and epiblast stem cells"

Fig.S1

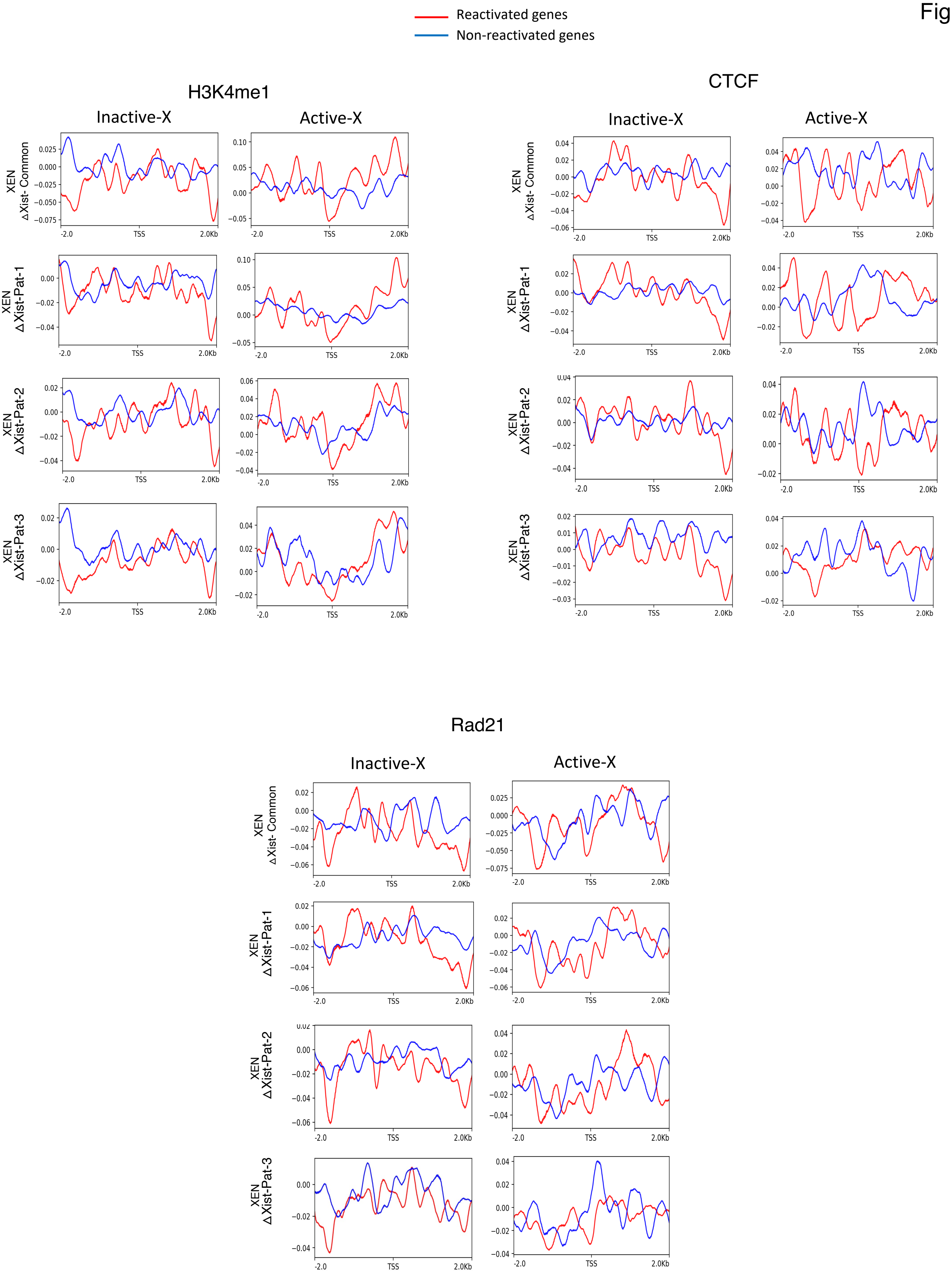

Reactivated genes
Non-reactivated genes
H3K4me1
CTCF
Rad21
Inactive-X
Active-X
XEN ΔXist- Common
XEN ΔXist-Pat-1
XEN ΔXist-Pat-2
XEN ΔXist-Pat-3
-2.0
TSS
2.0Kb

**Figure S1: Profiling of chromatin states of reactivated vs. non-reactivated genes.** Plots showing the enrichment of H3K4me1, CTCF, and Rad21 at TSS (+/- 2Kb) of non-reactivated (blue) and reactivated genes (red) on the inactive and active X chromosome.

Fig.S2

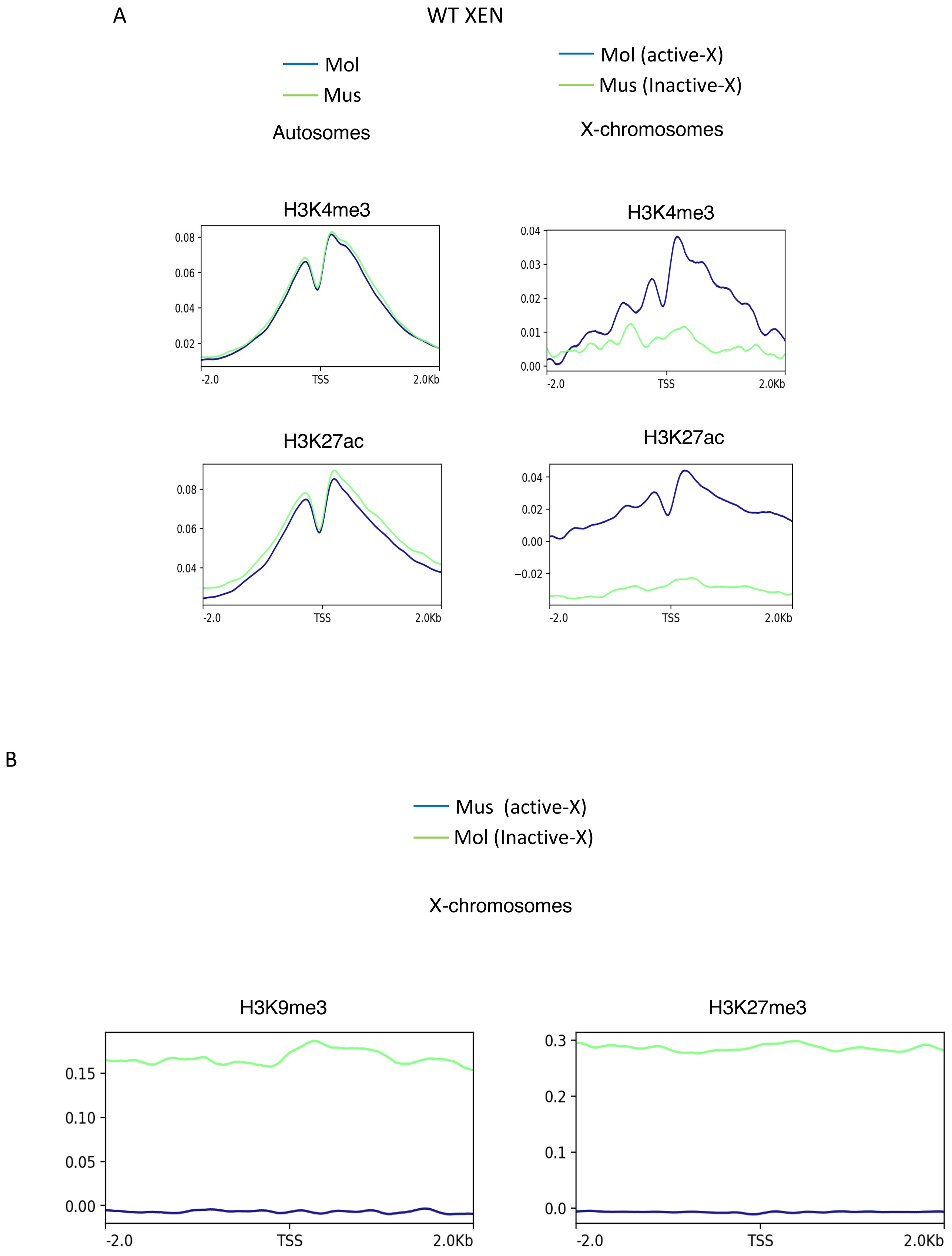

A
WT XEN
Mol
Mus
Autosomes
Mol (active-X)
Mus (Inactive-X)
X-chromosomes
H3K4me3
0.08
0.06
0.04
0.02
-2.0
TSS
2.0Kb
H3K4me3
0.04
0.03
0.02
0.01
0.00
-2.0
TSS
2.0Kb
H3K27ac
0.08
0.06
0.04
-2.0
TSS
2.0Kb
H3K27ac
0.04
0.02
0.00
−0.02
-2.0
TSS
2.0Kb
B
Mus (active-X)
Mol (Inactive-X)
X-chromosomes
H3K9me3
0.15
0.10
0.05
0.00
-2.0
TSS
2.0Kb
H3K27me3
0.3
0.2
0.1
0.0
-2.0
TSS
2.0Kb

**Figure S2: Profiling of allelic enrichment of chromatin marks in autosomes and X-chromosomes.** (A) Plots showing the enrichment of H3K4me3 and H3K27me3 at TSS (+/- 2Kb) of Mol (blue) and Mus (red) allele of autosomes and X-chromosome in WT XEN cells (B) Plots showing the enrichment of H3K9me3 and H3K27me3 at TSS (+/- 2Kb) of Mol (blue) and Mus (red) allele of autosomes and X-chromosome in EpiSC.
